# Arp2/3-dependent actin assembly shapes endosomes and promotes intracellular trafficking in fission yeast

**DOI:** 10.1101/2024.03.28.587130

**Authors:** Alejandro Melero, Olivia Muriel-Lopez, Miguel Basante-Bedoya, Sophie G Martin

## Abstract

Endosomes serve as crucial sorting centres that streamline the distribution of cell surface proteins. The early endosome receives traffic from both the plasma membrane (PM) and the Golgi, and orchestrates the redistribution of cargoes for recycling to the PM or through retrograde movement to the Golgi, and for degradation to late endosomes and lysosomes ^1^. In animal cells and amoebae, Arp2/3-mediated F-actin assembly plays critical roles in many aspects of endosome function, from capturing cargo to generating the forces required for membrane deformation and creating cargo-enriched transport carriers, thus promoting both recycling and degradative trafficking routes ^2^. Yeast models, which allowed dissection of the major membrane trafficking routes ^3^, exhibit highly simplified endosomes, as shown in *Saccharomyces cerevisiae*, where the trans-Golgi network (TGN) functions as recycling endosome ^4^. Furthermore, there is no reported role for Arp2/3 or F-actin in endomembrane remodelling; in fact, the Arp2/3 activators on animal endosomes – the WASH complex promoting recycling of transmembrane receptors towards the PM or TGN ^5,6^, and Annexin A2 necessary for endosome maturation and the movement of cargoes towards the degradative pathway ^7^ – do not exist in yeast cells ^8,9^. Here, we report that Arp2/3 and F-actin are present on fission yeast endosomes. Through live-imaging and correlative light-electron tomography, we demonstrate that Arp2/3 activity controls endosomal morphology and that this activity is essential for trafficking from the endosome to the degradative vacuole. Thus, Arp2/3-dependent actin assembly has a deeply conserved role in shaping and promoting the function of the endomembrane trafficking system.

## Results and discussion

In yeast cells, only three actin structures are thought to exist during mitotic proliferation, each well-described, well-insulated from the others, and assembled by a dedicated nucleating factor ^10,11^: Arp2/3, activated by WASP and type-I myosin, nucleates branched networks forming actin patches essential for the formation of endocytic vesicles at the PM ^12^; in *Schizosaccharomyces pombe*, the yeast we used in this study, two distinct formins, Cdc12 and For3, assemble the cytokinetic division ring and long actin cables in the cytosol, respectively. A third formin Fus1 assembles an actin fusion structure during sexual reproduction ^13^. We recently discovered an uncharacterized role for the Arp2/3 complex beyond its established roles in clathrin-mediated endocytosis. Indeed, we reported that inhibition of Arp2/3 activity leads to the reversible accumulation of sterols in an endomembrane compartment marked by the Soluble *N*-ethylmaleimide-Sensitive Factor Attachment Protein Receptor (SNARE) Syb1 (Synaptobrevin), indicating a novel function for branched actin networks in sterol trafficking in fission yeast and suggesting a possible role of Arp2/3 on endosomes ^14^.

To test whether Arp2/3 modulates endosome function, we repeated experiments of Arp2/3 inhibition with CK666, monitoring endosomal morphology with GFP-Syb1 and sterol-rich membranes using the sterol biosensor mCherry-D4H ^14^, for over 60 min and after Arp2/3 re-activation upon CK666 wash-out (Video 1). As previously described ^14^, Arp2/3 inhibition led to sterol accumulation on Syb1-marked endosomes, with internal D4H signals visible 20-30 min after CK666 addition (Fig 1A; magenta arrowheads). Concomitant with the internal signal increase, measurement of mCherry-D4H fluorescence intensity at the PM showed a strong drop from 20-30 min, reaching its lowest level after 40 min, and recovering fully within 20 min of CK666 washout (Fig 1B). Due to endocytic block, the GFP-Syb1 SNARE, which cycles between PM and early endosomes was partly trapped at the PM upon Arp2/3 inhibition. Interestingly, Arp2/3 inhibition had a rapid impact on endosome morphology, causing them to shift from elongated structures to more spherical objects within 10 min (Fig. 1A; green arrowheads). To quantitatively assess these morphological changes, we captured Airyscan super-resolution confocal stacks of GFP-Syb1 every min after CK666 addition and used automated segmentation techniques to obtain 3D objects and measure their volume. Strikingly, the volume of GFP-Syb1 labelled endosomes was significantly reduced within the first 10 min of CK666 addition (Fig 1C). This is also visible in the 3D rendering of example endosomes (Fig 1D, Fig S1A). Arp2/3 reactivation upon CK666 wash-out similarly caused fast endosome morphology changes within the first 10 min, with increases in volume (Fig. 1C), and the development of irregular morphologies when visualized in 3D (Fig 1D, Fig S1A). Thus, changes in Arp2/3 activity have rapid consequence on endosome morphology in fission yeast.

**Figure 1:**
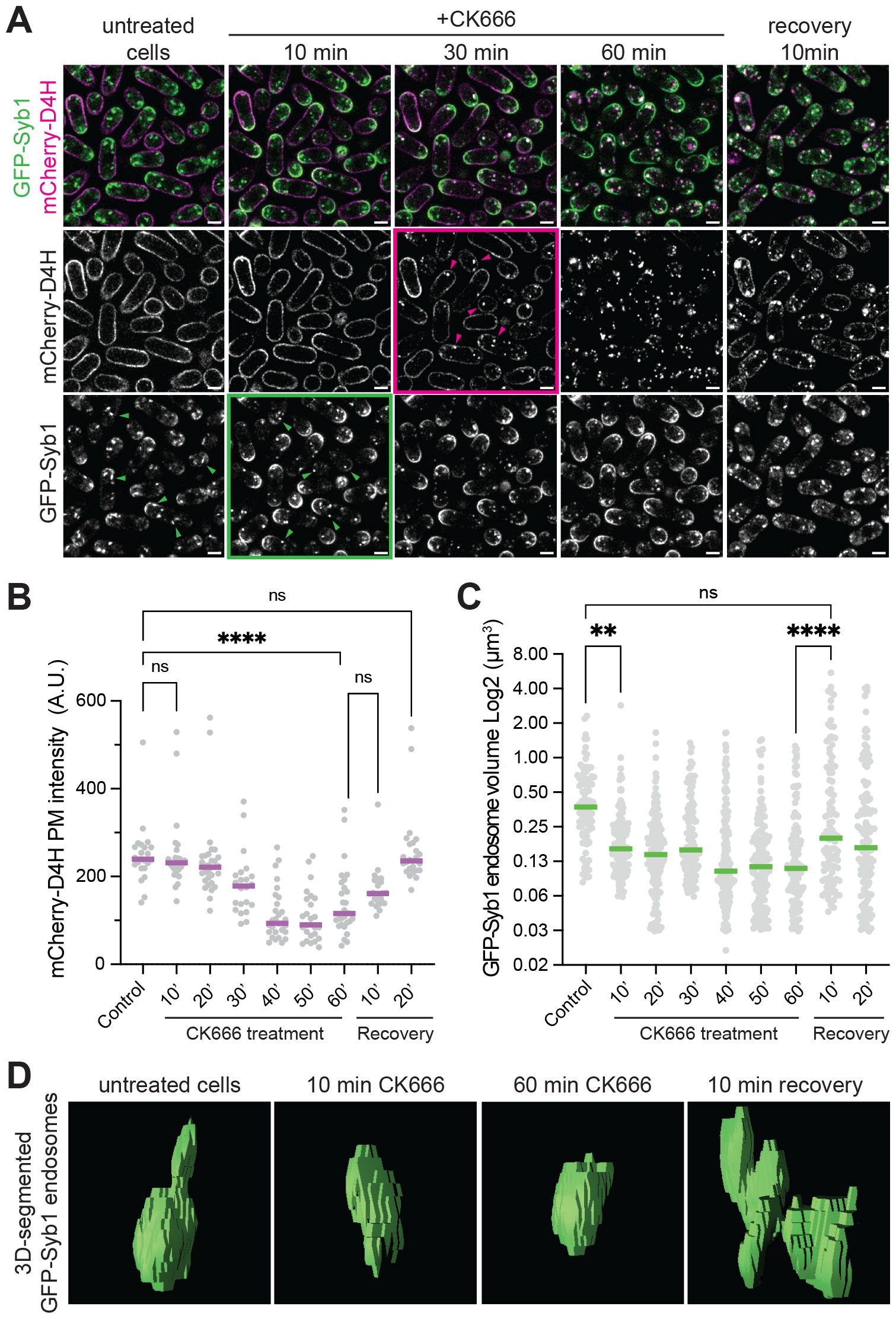
Arp2/3 activity modifies endosome morphology. **A)** Cells expressing the GFP-Syb1 endosomal marker and the mCherry-D4H sterol biosensor incubated with CK666 for 60 min, washed and followed for recovery for 10 min. Note the elongated shape of endosomes in untreated cells (green arrowheads). Magenta and green boxes indicate key timepoints after CK666 addition for mCherry-D4H internalization (30 min; magenta arrowheads) and endosome morphology remodelling (10 min; green arrowheads point to round endosomes), respectively. Scale bar = 2 μm. **B)** mCherry-D4H PM fluorescence signal in cells as in (A). Dots show individual cells’ median cortical signal; magenta line shows population median (n=31 cells). **C)** Endosome volumes obtained by supervised automated segmentations of GFP-Syb1 endosomes in cells as in (A). Dots show individual endosomes’ volume (μm^3^); green line shows population median (n=120 endosomes in 21 cells). **D)** Supervised automated segmentations of GFP-Syb1 endosomes at indicated timepoints. ANOVA was used as statistical test; ** p-value < 0.002; **** p-value < 0.0001.

While Arp2/3 inhibition affects both endosome morphology and sterol distribution, our time lapse imaging suggests that changes in endosome morphology precede sterol re-distribution. To more stringently test whether endosome morphological changes occur independently of sterol internalization, we used a *ltc1Δ* mutant strain, in which sterols remain at the plasma membrane even upon CK666 treatment ^14^. Ltc1 is a lipid transporter, required for the non-vesicular retrograde movement of sterol from the plasma membrane ^14^. In *ltc1Δ* cells, Arp2/3 inhibition with CK666 caused endosome morphology changes as in wild type cells, although sterol levels at the plasma membrane where unaffected (Fig. S1B). Thus, changes in endosome morphology after CK666 treatment are not a consequence of sterol re-routing towards endosomes but are a more proximal consequence of Arp2/3 inhibition.

To investigate endosome morphology at the ultrastructural level, we employed electron tomography and a correlative light-electron microscopy (CLEM) approach to specifically target GFP-Syb1 and mCherry-D4H labelled organelles. In short, the cells were cryo-immobilized using high pressure freezing and then embedded in Lowicryl polymer by freeze substitution. This method preserves both the fine ultrastructure and protein fluorescence ^15^, allowing us to obtain ultrastructural data on Syb1-labelled organelles through alignment of light and electron microscopy images (Fig 2A). We first examined the native ultrastructure of early endosomes in untreated *S. pombe* cells: wild type endosomes appeared as tubulo-cisternal structures of varied length (Fig 2A, Fig S2A), exhibiting frequent fenestrations and a tube diameter between 25 to 75 nm (Fig 2E). We note that the rare cytosolic signals of mCherry-D4H, labelling sterol-rich membranes, correlated with small vesicles (Fig 2A), suggesting that sterol can also move between organelles through vesicular pathways in fission yeast.

**Figure 2:**
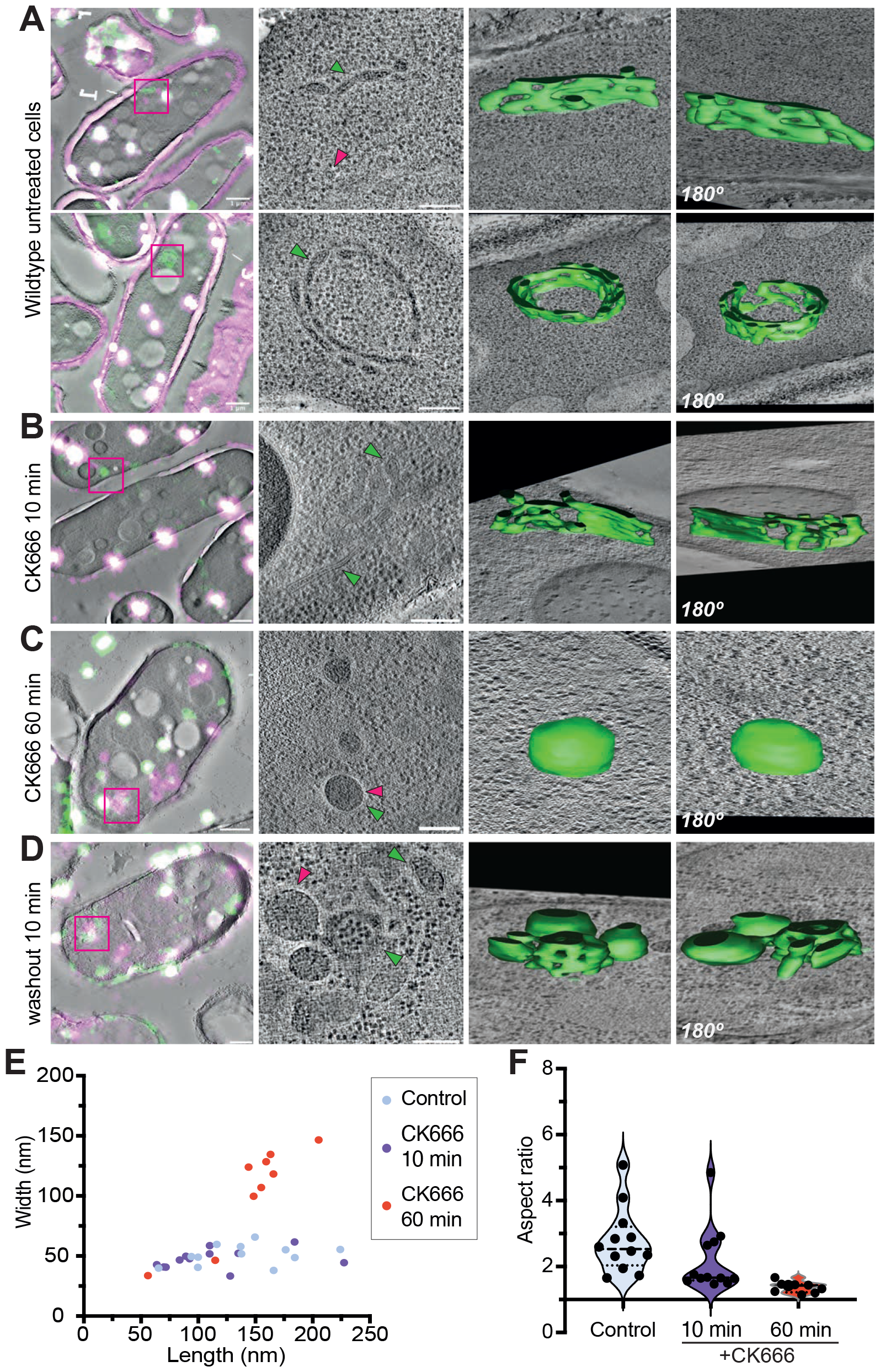
Ultrastructural description of endosomes in fission yeast cells. **A-D)** Correlated GFP-Syb1 (green) and mCherry-D4H (magenta) signal on electron tomogram virtual slice in (A) control cells, (B) cells treated with CK666 for 10 min, (C) cells treated with CK666 for 60 min and (D) cells recovering from CK666 treatment for 10 min. From left to right: Left, correlated overlay of light and electron microscopy (scale bar 1 μm). The white dots are fiducial tetraspeck beads used for image alignment. Central, high magnification virtual slice across the tomographic stack highlighting endosome ultrastructure (scale bar 200 nm). Arrowhead point to GFP-Syb1 (green) and mCherry-D4H (magenta) structures. In (A), a second mCherry-D4H-labelled vesicle is out of the range shown. Right panels, two side views of segmentation models of endosome ultrastructure. **E-F)** Morphometric analysis of endosomes in control and CK666-treated cells for 10 and 60 min, showing (E) width and length and (F) aspect ratio of endosomal cisterna measured every 20 virtual slices. Dots show the median values of X and Y axis for each endosomal cisterna. In (F), dashed lines show population medians; dotted lines indicate quartiles. n=12 in control and CK666 10 min, n=9 in CK666 60 min.

To observe the changes in endosome ultrastructure upon Arp2/3 inactivation and re-activation, we processed yeast cells for CLEM 10 and 60 min after CK666 addition, as well as 10 min after CK666 removal. After 10 min CK666, Syb1-labelled endosomes had reduced cisterna lengths and extended budding profiles (Fig 2B and E, Fig S2B and Fig S3A). After 60 min CK666 treatment, the ultrastructure of endosomes was altered into large ellipsoids lacking fenestrations or tubules, with diameters exceeding 100 nm (Fig 2C and E, Fig S2C and Fig S3B) ^14^. Rounding of endosomes was also visible in x-y aspect ratio measurements of cisterna, which rapidly reduced from >2.5 to ∼1 (Fig 2F). The larger x-y dimensions of endosomes after 60 min CK666 treatment (Fig 2E) likely correspond to shorter z dimensions, as many endosomes are fully contained in the tomogram, but not those of earlier timepoint or untreated samples. 10 min after CK666 removal, most ellipsoidal structures remodelled into fenestrated tubulo-reticular structures attached to large ellipsoidal vesicles (Fig 2D, Fig S2D, Fig S3C). Moreover, these endosomes exhibited multiple constriction sites and small budding profiles (Fig S3C), suggestive of active vesicle formation from the endosomal surface, and thus endosomal trafficking activity. These observations indicate that Arp2/3-dependent F-actin assembly is necessary to sustain the native tubulo-reticular ultrastructure of endosomes, which may in turn facilitate vesicular trafficking.

Neither Arp2/3 nor branched F-actin have been unambiguously identified at yeast endosomes, which may be partially due to the abundance of actin filaments that interact with or are coincidentally located near endosomes (Fig 3A). To facilitate the observation of branched actin networks, we took advantage of a *for3Δ* strain, which lacks the linear actin cables spanning the yeast cytoplasm ^16^. In *for3Δ* cells, F-actin structures labelled with Lifeact-mCherry can be found in close proximity to GFP-Syb1 endosomes (Fig 3B, Fig S4A). Using high frame rate Airyscan super-resolution microscopy with sub-second resolution, we observed fast morphological changes in endosomes, likely reflecting changing tube morphology, tube extension and/or connection (Fig 3C). Lifeact-mCherry formed patch-like structures associated with endosomes. These patches were not restricted to the cell cortex (Fig S4A), but were also found in the cell interior and often coincided with the tips and sites of constrictions of the endosomal tubular network (Fig 3B-C, arrowheads). We also co-imaged Arc5-mCherry, a subunit of the Arp2/3 complex, and GFP-Syb1 (Fig 3D, Fig S4B), which showed similar dynamics of small Arc5 foci following the reorganization of the endosomal network (Fig 3E, Fig S4B). These coincident signals with endosomes may in part stem from incoming actin-coated endocytic vesicles, which travel some distance in the cell interior towards endosomal compartments in both *S. pombe* and *S. cerevisiae* ^17,18^. However, the association of branched F-actin with endosomes over several seconds and its role in shaping these organelles suggest a proximal role in endosome remodelling beyond passive disassembly from endocytic vesicles.

**Figure 3:**
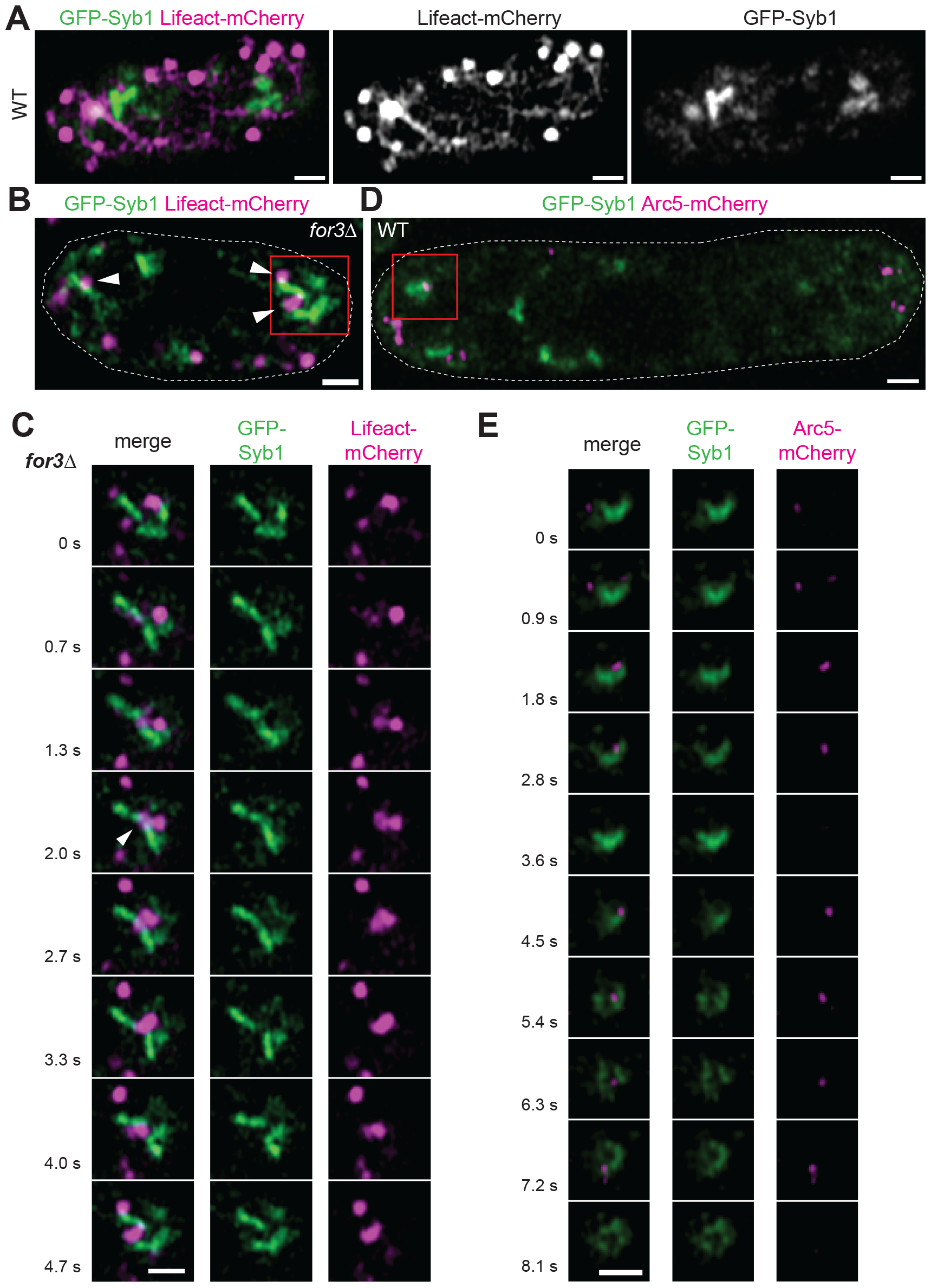
F-actin and Arp2/3 are present on endosomes. **A)** WT cell expressing GFP-Syb1 and Lifeact-mCherry. **B)** *for3Δ* cell expressing GFP-Syb1 and Lifeact-mCherry showing several Lifeact-mCherry foci in contact with the endosomal network. Arrowheads mark cell-internal actin structures in contact with endosomes; square highlights timelapse montage in (C). **C)** Time-lapse sequence of the square in (B). Arrowhead points to one of the Lifeact-mCherry signals at a site of endosome remodelling. **D)** WT cell expressing GFP-Syb1 and Arc5-mCherry showing an example of Arc5-mCherry foci on the endosomal network. Square highlights timelapse montage in (E). **E)** Time-lapse sequence of the square in (D), showing persistent presence of Arc5 on the endosome as it changes morphology. Scale bars = 1 μm.

To understand the functional role of branched F-actin on endosomal trafficking and probe its possible role in facilitating trafficking away from the endosome, we followed the internalization of the membrane dye FM4-64 from the plasma membrane to the vacuole ^18,19^. In control conditions, FM4-64 appeared at early endosomes marked with GFP-Syb1 in the first 5 min after addition, and further trafficked towards vacuoles within 20 to 30 min (Fig 4A, S5A). After 30 min incubation, most FM4-64 signal was localized at vacuoles, and only a few endosomes remained labelled with the dye. To observe the effect of Arp2/3 inhibition for endosomal trafficking, we incubated yeast cells with FM4-64 for 4 min, subsequently washing the excess dye with media containing CK666, thus effectively blocking Arp2/3 function upon FM4-64 reaching early endosomes. Remarkably, upon Arp2/3 inhibition, FM4-64 did not reach the vacuole after 30 min of incubation but remained associated with endosomal structures (Fig 4B). Thus, trafficking from the endosome to the degradative route requires Arp2/3 activity. Recycling of FM4-64 to the PM was also attenuated in the presence of CK666, but we note that some trafficking to the PM may proceed in absence of Arp2/3 activity, as judged from the accumulation of Syb1 at the PM. At the 30 min time point, Arp2/3 reactivation by CK666 wash-out reactivated the transport of FM4-64 towards the vacuoles, which were already clearly observable within 5 min and strongly labelled after 20 min (Fig 4C). A few endosomes remained labelled by FM4-64 20 min after CK666 wash-out, similar to late timepoints in control conditions. The fast arrival of FM4-64 in vacuoles after CK666 washout suggests that some aspects of endosome maturation proceed normally without Arp2/3 function but transfer to vacuoles is impeded. We conclude that Arp2/3 and branched F-actin are required downstream of endocytosis for membrane trafficking, likely by promoting membrane remodelling and the formation of vesicles away from the endosome.

**Figure 4:**
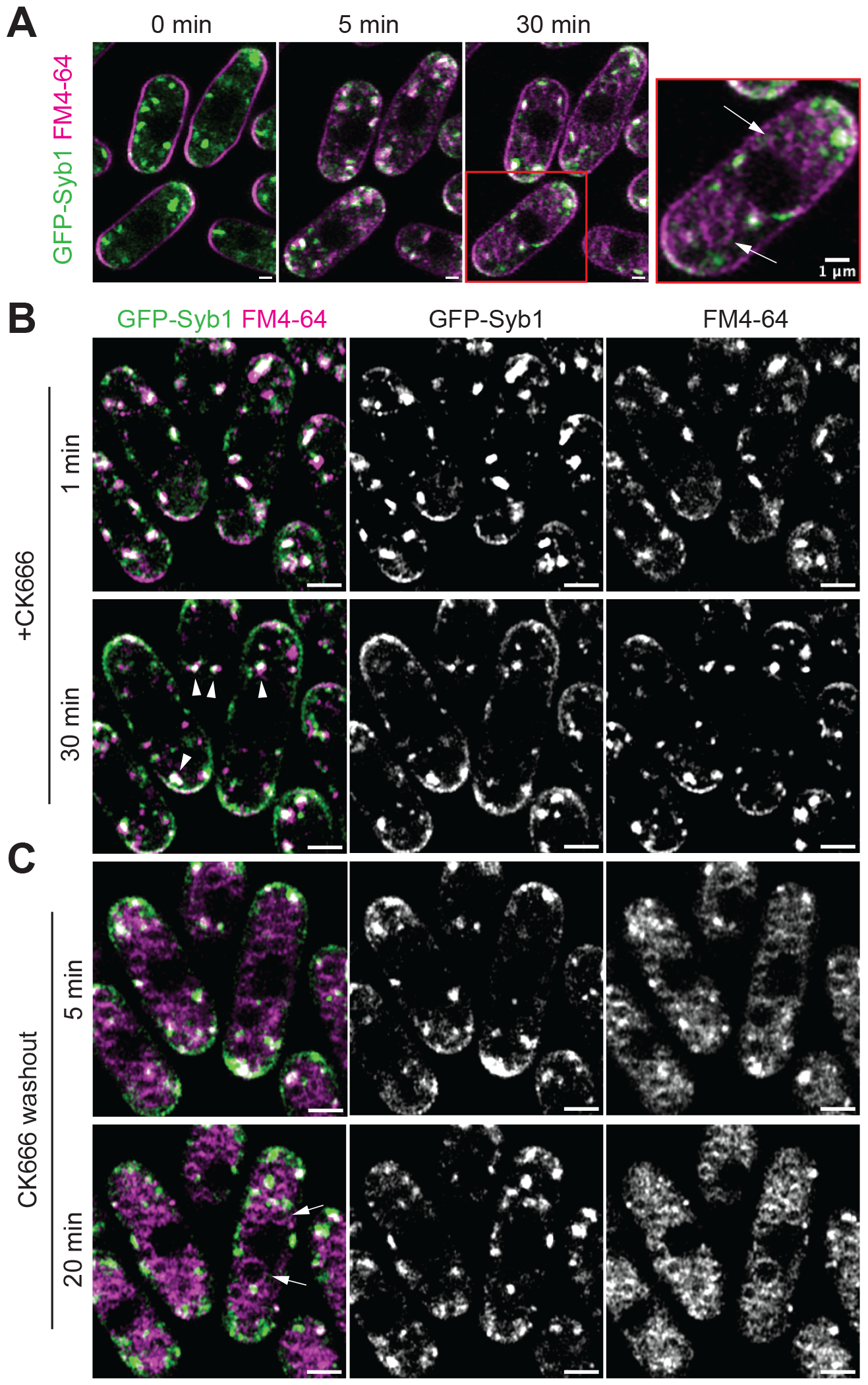
Arp2/3 activity is required for trafficking from the endosome to the vacuole. **A)** Timing of internalization of the membrane dye FM4-64 in GFP-Syb1-expressing cells. Right panel: highlight on vacuolar FM4-64 accumulation (arrows) after 30 min of incubation. **B)** GFP-Syb1-expressing cells incubated with FM4-64 for 4 min, washed with CK666-containing media and imaged 1 and 30 min post CK666 addition (5 and 34 min post FM4-64 addition). The progression of the dye towards vacuoles is interrupted upon addition of CK666, even after 30 min. Arrowheads point to a few FM4-64-labelled endosomes. **C)** Same group of cells as in (B), after washing out CK666. FM4-64 immediately progresses to the vacuoles (arrows). Scale bars 1 μm (A) and 2 μm (B, C).

## Conclusion

We have shown that Arp2/3 complex associates with early endosomes in fission yeast, promoting the formation of branched F-actin structures that play a critical role in shaping the organelle and supporting its trafficking function towards the vacuole. Our ultrastructural data suggest that Arp2/3 activity facilitates the transformation of endosomes into fenestrated tubulo-cisternal networks. The tubulo-cisternal nature of early endosomes has been described across organisms by early ultrastructural studies ^20-22^ and more modern approaches of whole cell block-face CLEM ^23^. To date, no specific protein responsible for conferring this tubular structure has been identified for endosomes, unlike the reticulons found at the ER ^24^ and the golgins at the Golgi complex ^25^. The narrow tube diameters observed in early endosomes, both in mammalian cells (<60 nm, ^20,26^) and in our yeast studies (∼50 nm), do not form spontaneously but necessitate the application of active forces for their generation and maintenance. Thus, a likely important function of F-actin on endosomes is to provide force for membrane deformation.

The high curvature radius of endosomal networks may in turn facilitate coat binding and the budding of trafficking intermediates. For instance, recent research has revealed that high curvature is a prerequisite for retromer to effectively enforce membrane curvature ^27^. A pre-requisite for high curvature to facilitate budding is common in other organelles for various coats, including COPI ^28^ and COPII ^29,30^. Our data show that Arp2/3 activity is required for trafficking from endosome to vacuole in fission yeast. This is similar to its role in endosome maturation in mammalian cells ^31^, even though neither of the mammalian activators – Annexin A2, which promotes endosome maturation ^7^, and the WASH complex, which promotes not only cargo recycling ^5,6^, but also the transport of cargo towards the late endosome ^32^ – are conserved in fission yeast ^8,9^. Thus, an open question is how the Arp2/3 complex is recruited and activated on endosomes in fission yeast. Other nucleation promoting factors, of which fission yeast express several may promote Arp2/3 activity on endosomes, although we could not be convinced of the localization of either Wsp1, Myo1, Abp1, Dip1 or Crn1 to Syb1-labelled endosomes. Alternatively, branched actin assembly on early endosomes may be seeded by other actin structures, for instance the actin coat that powers the motility of incoming endocytic vesicles ^33,34^, or actin cables on which these endocytic vesicles travel ^17,18^. Given the simplicity of the yeast endomembrane system and that a late Golgi compartment serves as recycling endosome ^4^, the molecular pathway of F-actin assembly on yeast endosomes may bear similarities to those functioning at the TGN in animal cells, where carrier vesicle formation for trafficking to the lysosome requires clathrin and actin-assembly ^35,36^. Identifying the mechanisms of F-actin assembly on endosomes, and their similarities and differences from the well-established mechanisms of clathrin-mediated endocytosis at the PM, are important challenges for the future.

## Acknowledgements

We thank Christelle Genoud and Jean Daraspe (University of Lausanne) for help with electron microscopy, Magdalena Marek (Max Planck Genome Centre Cologne) for the construction of several strains, and Fred Chang (UCSF) for the Arc5-mCherry strain. This work was funded by a Swiss National Science Foundation (310030_191990) and ERC CoG grant (CellFusion) to SGM.

## Author contributions

AM and SGM designed the project. AM conducted all experiments, with help from OM for the preparation, acquisition and correlation of CK666 recovery tomograms, and from MBB for the acquisition of the *for3Δ* Lifeact-mCherry GFP-Syb1 timelapses in Figures 3B-C and S4A. AM performed all image analysis and quantifications. AM and SGM wrote the paper. SGM provided supervision and acquired funding.

## Materials and methods

### Contact for reagent and resource sharing

Further information and requests for resources and reagents should be directed to and will be fulfilled by Sophie G. Martin.

### Experimental Model and growth conditions

*Schizosaccharomyces pombe* strains were grown in Edinburgh minimal medium supplemented with amino acids (EMM-ALU) or in rich YE5S at 25°C. Standard genetic methods and growth conditions were used. Strains are listed in Table S1.

**Table S1:**
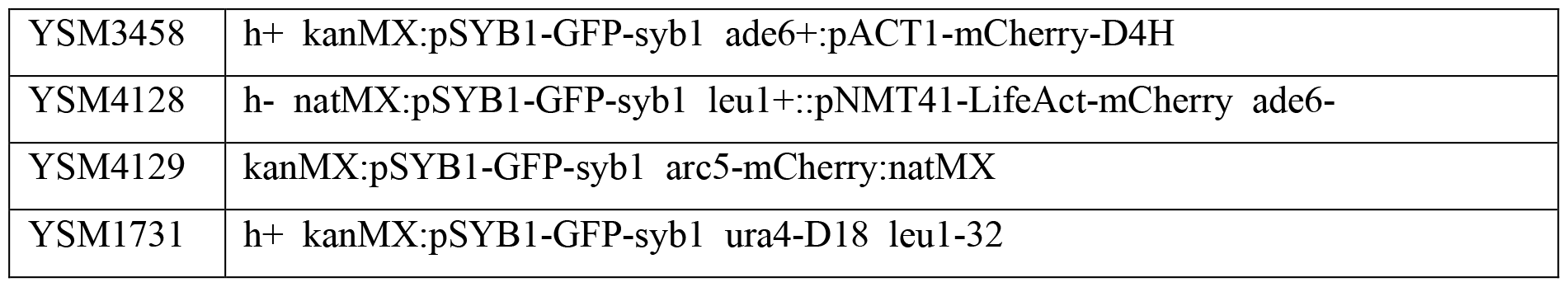
Strains used in this study.

To block Arp2/3 complex CK666 at a final concentration of 500 μM was used from a 50 mM stock solution in DMSO. For recovery from CK666 treatment, cells incubated with CK666 for 1 h were washed ten times with medium. For visualization of FM4-64 uptake, cells were stained with 5 μM FM4-64 dye for 4-5 min in EMM-ALU medium. The dye was washed out five times with EMM-ALU with or without CK666 upon reaching early endosomes.

### Microscopy

#### Live imaging sample preparation

Yeast cells were imaged using Ibidi μ-Slide VI 0.5 Glass Bottom, (ref. 80606, IBIDI GMBH, Germany). The chambers were treated with soybean lectin (*Glycine max* lectin, Sigma-Aldrich) for 1h ahead of the experiment and washed with EMM-ALU before adding the cells. Yeasts were grown overnight on EMM-ALU up to 0.5-0.6 OD_600_/ml, 1 ml of culture was spun down, and cells were resuspended in 150 μl of fresh medium and inoculated in the chamber. After 5 min, the excess of non-adhered cells was washed away with medium, leaving a final volume of 150 μl in the chamber.

#### Imaging

All live microscopy images were acquired using a Zeiss LSM980 system with Plan-Apochromat 63x/1.40 oil differential interference contrast objective with the ZEN Blue software (Zeiss). Imaging was set in super-resolution SNR/Sensitivity mode mode with bidirectional scanning in either frame or line mode, as indicated below. All other settings were optimized as recommended by the ZEN Blue software.

#### CK666 treatment and recovery

Upon 5 min of recording, the acquisition was paused to add CK666, the medium was mixed in the chamber using a P200 micropipette and the acquisition restarted. After 60 min, the acquisition was paused and the media inside the chamber was exchanged 10 times with fresh EMM-ALU medium using a P200 micropipette. Then acquisition was re-started for 20 to 30 more min.

#### Plasma membrane mCherry-D4H signal intensity quantification

Quantification of plasma membrane signal intensity was done using FIJI software ^37^. At each indicated timepoint, the perimeter of each individual cell was selected manually using the segmented line tool and the median intensity signal quantified using the measurements tool. The data was represented and analysed applying a two-way ordinary ANOVA using GraphPad Prism statistical package.

#### GFP-Syb1 Endosome 3D imaging and volume segmentation

Cells were imaged in frame mode, acquiring one stack of 36 steps every 60 seconds with several yeast cells per field of view (pixel width 42.5 nm, voxel depth 170 nm). Stacks were processed with FIJI software using a 3D gaussian filter with a sigma of 1. Endosomes were automatically segmented using the FIJI plugin “3D Object Counter” ^38^ using default settings and manually curated using the “3D manager” plugin ^39^. Large clusters of GFP-Syb1 endosomes and cortical GFP-Syb1 were resolved using “split” tool, obtaining separated volume segmentations for the endosome and plasma membrane portion. The plasma membrane portion was discarded. Large endosome-plasma membrane clusters that could not be resolved were discarded and not used for quantification. Volume measurements and 3D rendering were obtained using the “3D manager options” and “3D manager”. Between 100 and 130 endosomes were analysed per time-point.

#### Fast super-resolution line scan of GFP-Syb1 with Lifeact-mCherry or Arc5-mCherry

A field of yeast cells was imaged in line scan mode at 42.5 nm pixel width and sub-second (two-colour) frame rate. To control for possible crosstalk between fluorophores a GFP-Syb1 only yeast strain was used as a control, adjusting the mCherry channel to avoid any unspecific detection of Lifeact-mCherry and Arc5-mCherry signals. Image series were processed with FIJI software using gaussian filter sigma 1.

#### CLEM

CLEM was as described ^40^. Cells were grown overnight in EMM-ALU at 25ºC to an OD_600_ of 0.5-0.6. 1.5 ml from the culture were gently centrifuged (3000 xg, 3 min) and the supernatant discarded to obtain a dense cell paste. The cell slurry was pipetted onto 3-mm-wide, 0.1-mm-deep specimen carriers (Wohlwend type A) closed with a flat lid (Wohlwend type B) for high-pressure freezing with a Wohlwend HPF Compact 02 (for control samples) or a Leica EM ICE (Leica Microsystems; rest of samples). The carrier sandwich was disassembled in liquid nitrogen before freeze substitution. High-pressure frozen samples were processed by freeze substitution using 1% uranyl acetate in acetone and embedding in Lowicryl HM20 using the Leica AFS 2 robot. 300-nm sections were cut with a diamond knife using a Leica Ultracut UC7 ultramicrotome, collected in 1x PBS, and picked up on carbon-coated 200-mesh copper grids (AGS160; Agar Scientific). Upon light microscopy, TetraSpeck™ Microspheres (Invitrogen, 0.1 μm) were used as CLEM fiducials. The microspheres were adsorbed onto the grids by blotting on a drop of 1x PBS with 1:200 dilution of beads for 10 min and three washes to discard excess of beads. For light microscopy, the grid was inverted onto a 1x PBS drop on a microscope coverslip, which was mounted onto a microscope slide and imaged as a z stack using the LSM980 in Airyscan SNR/Sensitivity mode. The grid was then recovered, rinsed in H_2_O, and dried before post-staining with Reynolds lead citrate for 10 min. 15-nm protein A–coupled gold beads were adsorbed to the top of the section as fiducials for tomography. Electron tomograms were acquired on a FEI Tecnai 12 at 120 kV using a bottom mount FEI Eagle camera (4kx4k). Low-magnification images were acquired at 17.816-nm pixel size and high magnification at 1.205-nm pixel size. For tomographic reconstruction of regions of interest, tilt series were acquired at 1.205-nm pixel size (Tecnai) over a tilt range as large as possible up to ±60° at 1° increments using the Serial EM software ^41^. The IMOD software package with manual or supervised automatic patch tracking alignment ^42,43^ was used for tomogram reconstruction. Segmentation and modelling were done on IMOD software as well. For light and electron microscopy correlation, the Icy software was used ^44^, using the plugin ec-CLEM ^45^.

#### Morphometric analysis of endosomes

Length and width of endosome cisterna cross-sections was done with FIJI software. The perimeter of each endosome section (cisterna) was selected using segmented lines and the Feret diameters were calculated using the measurements tool. This was done every 20 nm across the tomogram depth (∼200-300 nm). The median length and width, as well as the aspect ratio, were calculated on Excel for each endosome and plotted using Graphpad Prism 10.

---

**Figure S1:**
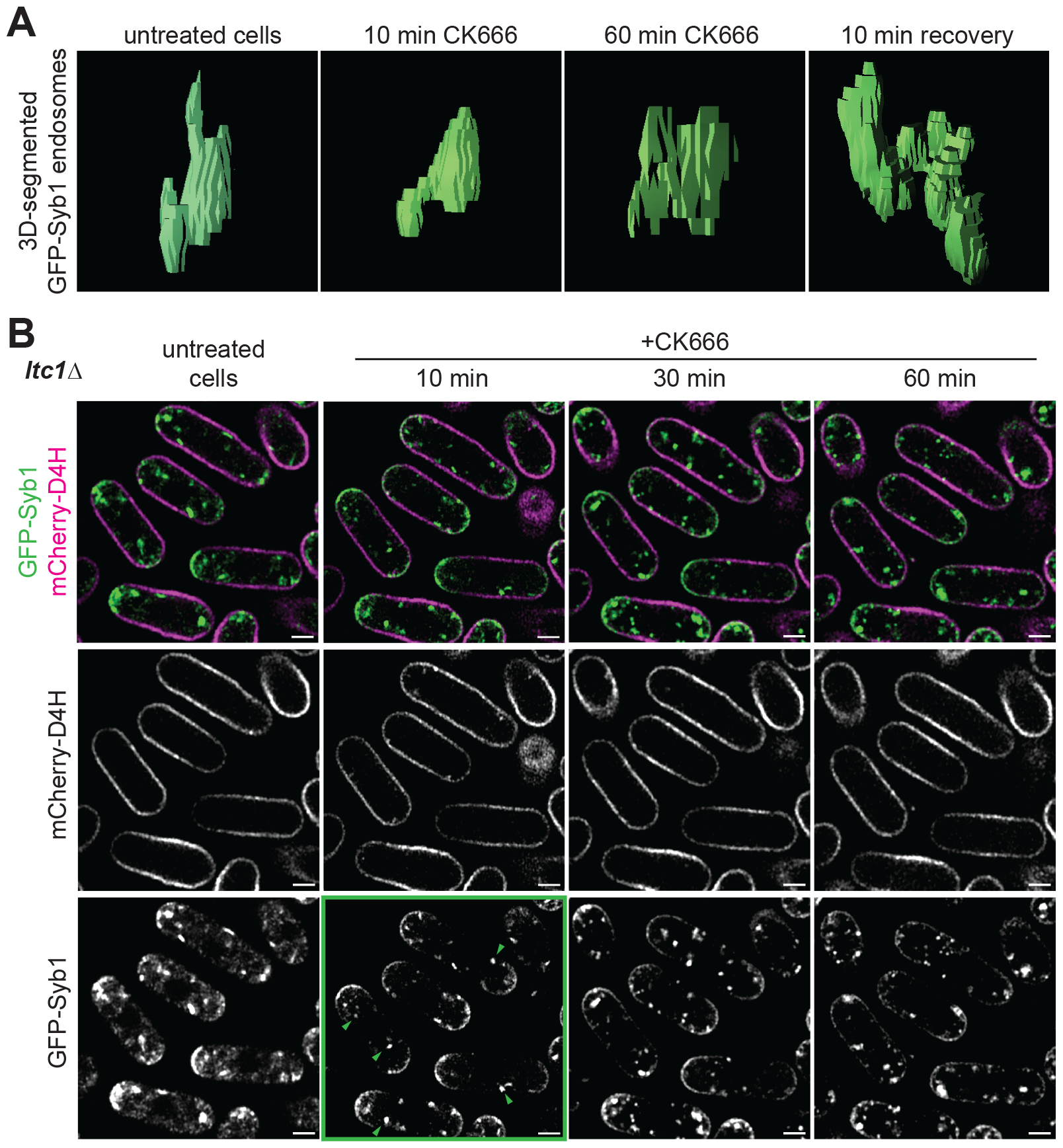
Endosome rounding after CK666 treatment is independent of sterol accumulation (related to Figure 1). **A)** Additional examples of supervised automated segmentations of GFP-Syb1 endosomes in WT cells at indicated timepoints, as in Fig 1D. **B)** *ltc1Δ* cells expressing GFP-Syb1 and mCherry-D4H incubated with CK666 for 60 min. Note that endosomes become round after 10 min (green highlight and arrowheads), even if sterol remains at the PM in this mutant. Scale bar = 2 μm.

**Figure S2:**
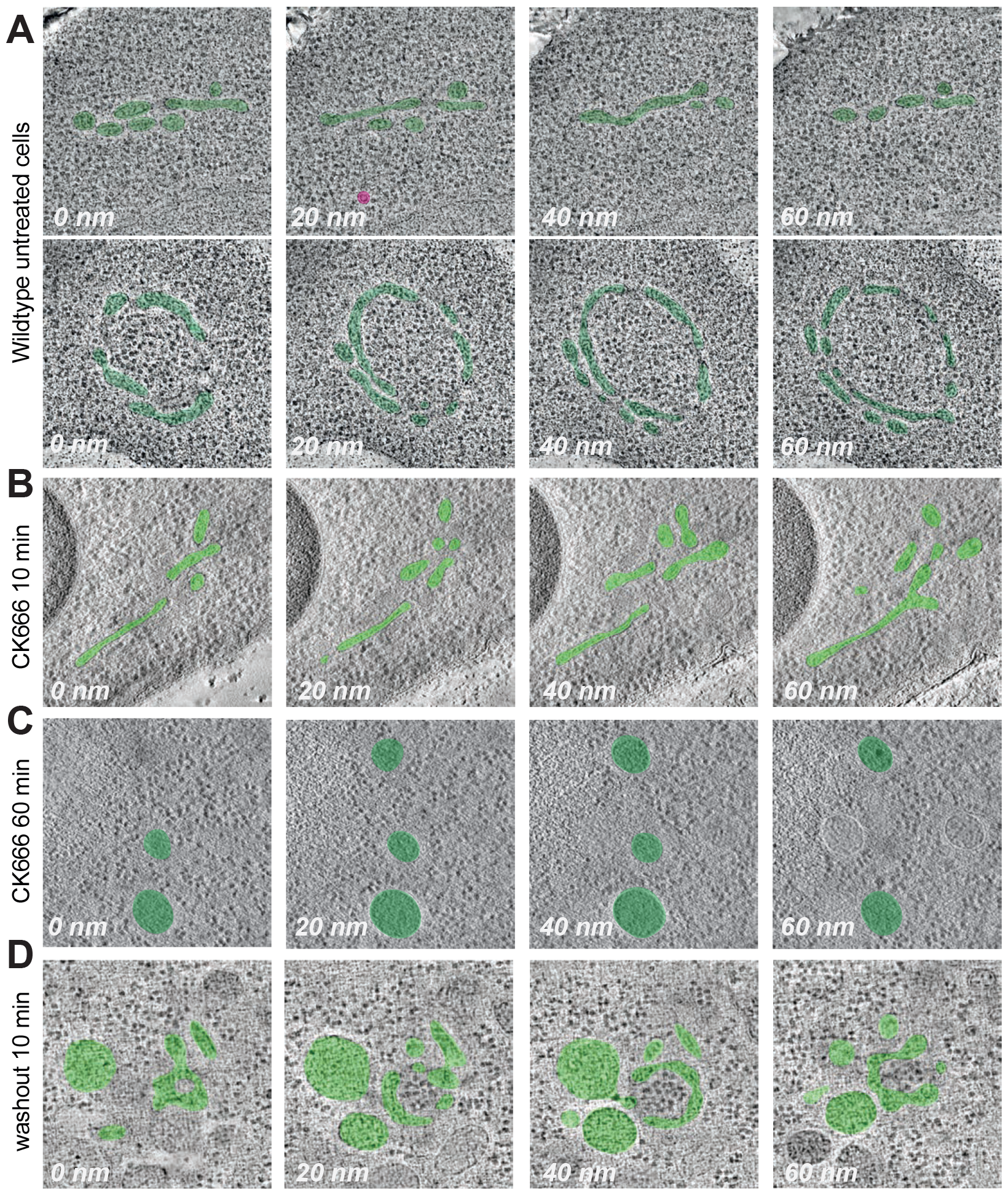
Ultrastructural description of endosomes in fission yeast cells (related to Figure 2). Virtual slices across the tomographic stack with false colour (green) highlighting the GFP-Syb1-labelled endosomes represented in Fig 2. **A)** Correlated GFP-Syb1 endosome in untreated cells from Fig 2A. A small vesicle corresponding to D4H signal is also shown (magenta; top row). A second mCherry-D4H vesicle is out of the range shown. **B)** Correlated GFP-Syb1 endosome in cells treated with CK666 for 10 min from Fig 2B. **C)** Correlated GFP-Syb1 endosome in cells treated with CK666 for 60 min from Fig. 2C. **D)** Correlated GFP-Syb1 endosome in cells 10 min after CK666 washout from Fig. 2D.

**Figure S3:**
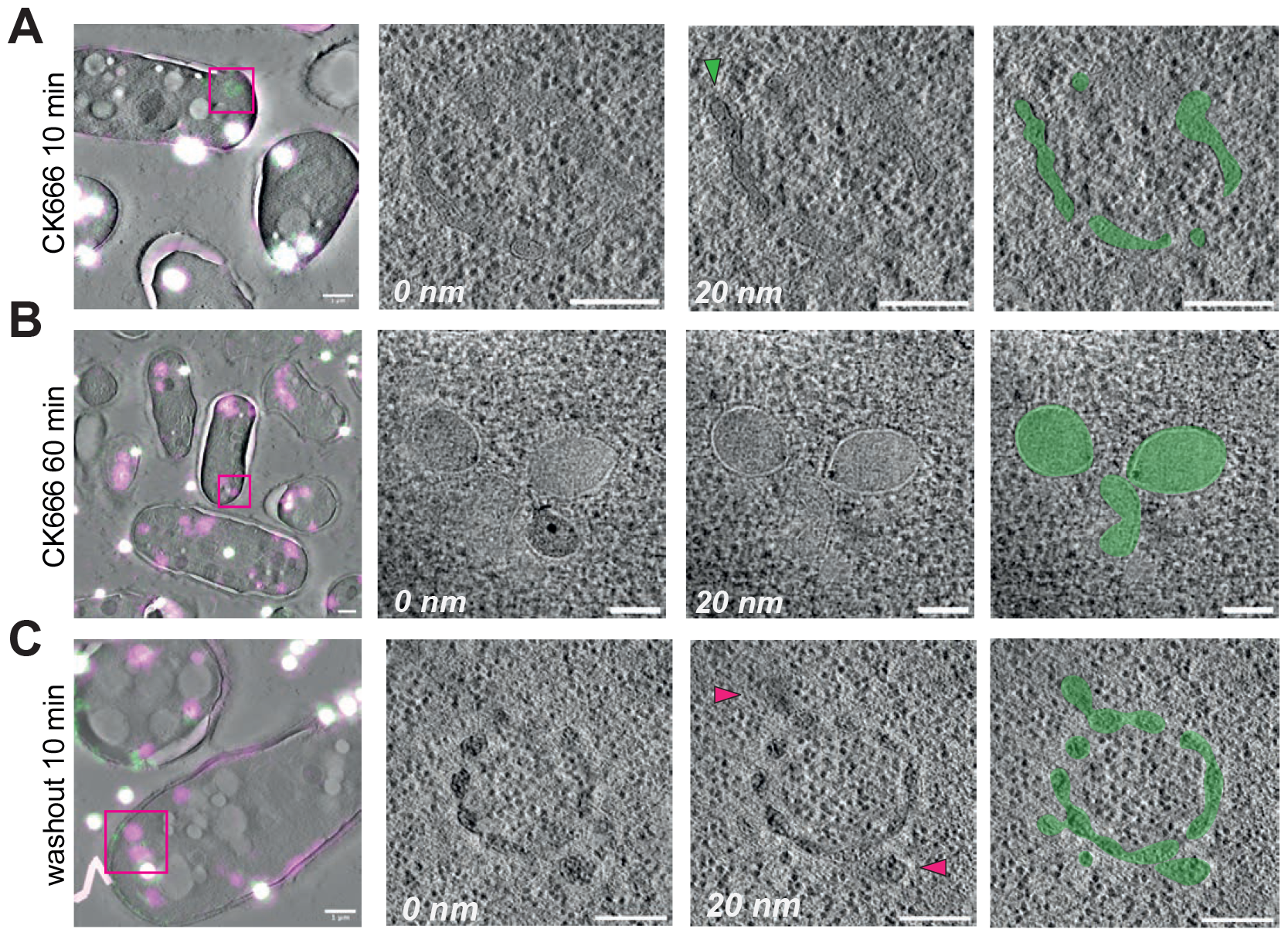
Ultrastructural description of endosomes in fission yeast cells (related to Figure 2). **A-C)** Additional examples of correlated GFP-Syb1 (green) and mCherry-D4H (magenta) signal on electron tomogram virtual slices in (A) cells treated with CK666 for 10 min, (B) cells treated with CK666 for 60 min and (C) cells recovering from CK666 treatment for 10 min. From left to right: Left, correlated overlay of light and electron microscopy (scale bar 1 μm). The white dots are fiducial tetraspeck beads used for image alignment. Middle, two high magnification virtual slices across the tomographic stack at 20 nm distance, highlighting endosome ultrastructure (scale bar 200 nm). Right panel, false colour highlighting the endosome. In (A), the green arrowhead highlights a coated vesicle budding event. In (C), the magenta arrowheads indicate areas of mCherry-D4H signal overlayed with the GFP-Syb1 signal.

**Figure S4:**
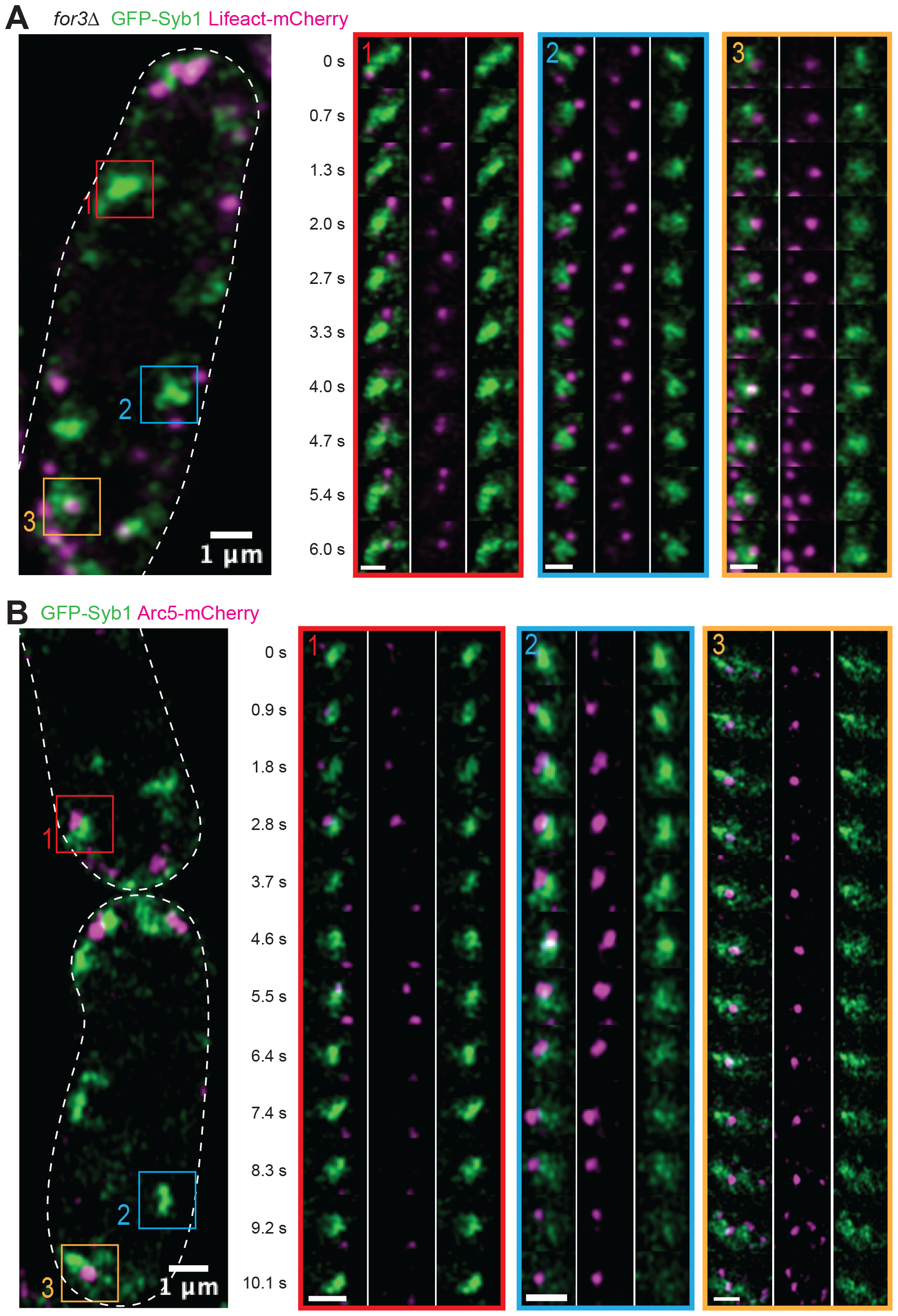
F-actin and Arp2/3 are present on endosomes (related to Figure 3). **A)** A *for3Δ* GFP-Syb1 Lifeact-mCherry cell highlighted with squares on three subregions with endosomes interacting with branched F-actin patches. Right panels correspond to timelapses of these endosome-branched actin interaction events. **B)** A GFP-Syb1 Arc5-mCherry cell highlighted with squares on three subregions with endosomes interacting with Arc5. Right panels correspond to time-lapses of these endosome-Arc5 interaction events. Scale bars = 1 μm.

**Figure S5:**
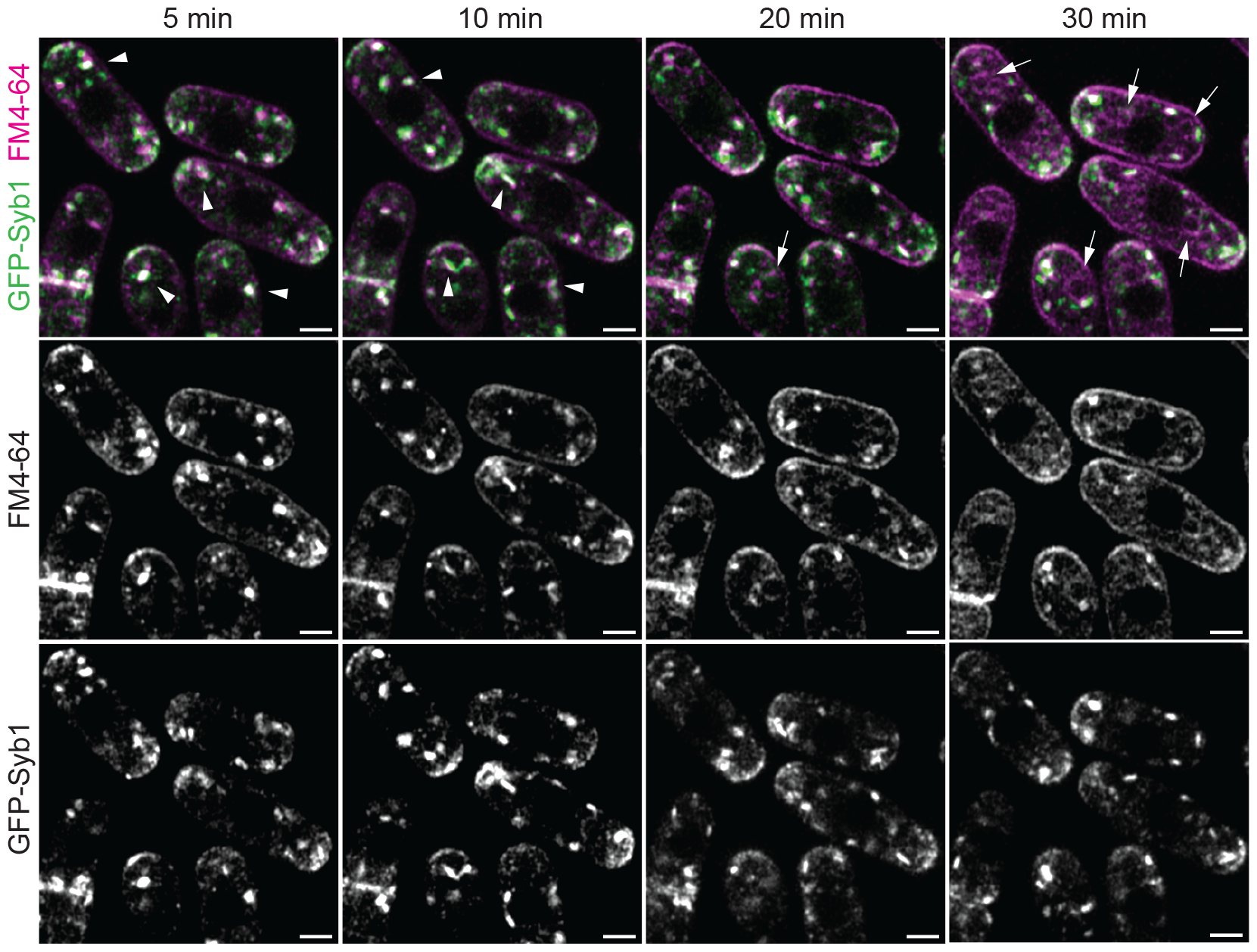
Time course of FM4-64 internalization in WT cells (related to Figure 4). Cells expressing GFP-Syb1 incubated with FM4-64 membrane dye and imaged at indicated time points, as in Fig 4A but with additional time points. FM4-64 labels Syb1-marked endosomes after 5 min (arrowheads). Weak vacuole staining can be detected after 20 min and stronger vacuole localization after 30 min (arrows). Scale bars = 2 μm.

**Video 1: Timelapse of endosome morphology changes upon Arp2/3 inhibition and re-activation**.

Cells expressing GFP-Syb1 and mCherry-D4H, imaged for 5 min before addition of CK666 for 60 min (+CK666; t = 0) and subsequent washout with fresh media (recovery; t = 0). Note the elongated shape of GFP-Syb1-labelled endosomal structure before treatment, change to round shape rapidly after CK666 addition before mCherry-D4H signal enrichment, and recovery upon washout. Scale bar = 3 μm.

**Video 2: Timelapse of F-actin and endosomes**.

Yeast cell expressing GFP-Syb1 and Lifeact-mCherry bearing a *for3Δ* deletion. Acquisition settings were such that a GFP-Syb1 strain without Lifeact-mCherry showed no magenta signal at all. Thus, even weak magenta signals are likely to be real Lifeact-mCherry signal. Time-lapse 0.7 seconds per frame. Scale bar = 2 μm.

**Video 3: Timelapse of Arc5 and endosomes**.

Yeast cell expressing GFP-Syb1 and Arc5-mCh. Acquisition settings were such that a GFP-Syb1 strain without Arc5-mCherry showed no magenta signal at all. Thus, even weak magenta signals are likely to be real Arc5 mCherry signal. Note that, in contrast to Lifeact-mCherry (video 2), most Arc5 signals are close to the cell cortex. Time-lapse 0.9 seconds per frame. Scale bar = 2 μm.

## References

1. Scott, C.C., Vacca, F., and Gruenberg, J. (2014). Endosome maturation, transport and functions. Semin Cell Dev Biol 31, 2–10. 10.1016/j.semcdb.2014.03.034.

2. Simonetti, B., and Cullen, P.J. (2019). Actin-dependent endosomal receptor recycling. Curr Opin Cell Biol 56, 22–33. 10.1016/j.ceb.2018.08.006.

3. Spang, A. (2015). Anniversary of the discovery of sec mutants by Novick and Schekman. Mol Biol Cell 26, 1783–1785. 10.1091/mbc.E14-11-1511.

4. Day, K.J., Casler, J.C., and Glick, B.S. (2018). Budding Yeast Has a Minimal Endomembrane System. Dev Cell 44, 56–72 e54. 10.1016/j.devcel.2017.12.014.

5. Gomez, T.S., and Billadeau, D.D. (2009). A FAM21-containing WASH complex regulates retromer-dependent sorting. Dev Cell 17, 699–711. 10.1016/j.devcel.2009.09.009.

6. Derivery, E., Sousa, C., Gautier, J.J., Lombard, B., Loew, D., and Gautreau, A. (2009). The Arp2/3 activator WASH controls the fission of endosomes through a large multiprotein complex. Dev Cell 17, 712–723. 10.1016/j.devcel.2009.09.010.

7. Morel, E., Parton, R.G., and Gruenberg, J. (2009). Annexin A2-dependent polymerization of actin mediates endosome biogenesis. Dev Cell 16, 445–457. 10.1016/j.devcel.2009.01.007.

8. Derivery, E., and Gautreau, A. (2010). Evolutionary conservation of the WASH complex, an actin polymerization machine involved in endosomal fission. Communicative & integrative biology 3, 227–230. 10.4161/cib.3.3.11185.

9. Braun, E.L., Kang, S., Nelson, M.A., and Natvig, D.O. (1998). Identification of the first fungal annexin: analysis of annexin gene duplications and implications for eukaryotic evolution. J Mol Evol 47, 531–543. 10.1007/pl00006409.

10. Mishra, M., Huang, J., and Balasubramanian, M.K. (2014). The yeast actin cytoskeleton. FEMS microbiology reviews 38, 213–227. 10.1111/1574-6976.12064.

11. Kovar, D.R., Sirotkin, V., and Lord, M. (2011). Three’s company: the fission yeast actin cytoskeleton. Trends Cell Biol 21, 177-187. S0962-8924(10)00257-6 [pii] 10.1016/j.tcb.2010.11.001.

12. Kaksonen, M., and Roux, A. (2018). Mechanisms of clathrin-mediated endocytosis. Nat Rev Mol Cell Biol 19, 313–326. 10.1038/nrm.2017.132.

13. Dudin, O., Bendezu, F.O., Groux, R., Laroche, T., Seitz, A., and Martin, S.G. (2015). A formin-nucleated actin aster concentrates cell wall hydrolases for cell fusion in fission yeast. J Cell Biol 208, 897–911. 10.1083/jcb.201411124.

14. Marek, M., Vincenzetti, V., and Martin, S.G. (2020). Sterol biosensor reveals LAM-family Ltc1-dependent sterol flow to endosomes upon Arp2/3 inhibition. J Cell Biol 219. 10.1083/jcb.202001147.

15. Kukulski, W., Schorb, M., Welsch, S., Picco, A., Kaksonen, M., and Briggs, J.A. (2011). Correlated fluorescence and 3D electron microscopy with high sensitivity and spatial precision. J Cell Biol 192, 111–119. 10.1083/jcb.201009037.

16. Feierbach, B., and Chang, F. (2001). Roles of the fission yeast formin for3p in cell polarity, actin cable formation and symmetric cell division. Curr Biol 11, 1656–1665.

17. Pelham, R.J., Jr., and Chang, F. (2001). Role of actin polymerization and actin cables in actin-patch movement in Schizosaccharomyces pombe. Nat Cell Biol 3, 235–244. 10.1038/35060020.

18. Huckaba, T.M., Gay, A.C., Pantalena, L.F., Yang, H.C., and Pon, L.A. (2004). Live cell imaging of the assembly, disassembly, and actin cable-dependent movement of endosomes and actin patches in the budding yeast, Saccharomyces cerevisiae. J Cell Biol 167, 519–530.

19. Vida, T.A., and Emr, S.D. (1995). A new vital stain for visualizing vacuolar membrane dynamics and endocytosis in yeast. J Cell Biol 128, 779–792. 10.1083/jcb.128.5.779.

20. Tooze, J., and Hollinshead, M. (1991). Tubular early endosomal networks in AtT20 and other cells. J Cell Biol 115, 635–653. 10.1083/jcb.115.3.635.

21. Stoorvogel, W., Oorschot, V., and Geuze, H.J. (1996). A novel class of clathrin-coated vesicles budding from endosomes. J Cell Biol 132, 21–33. 10.1083/jcb.132.1.21.

22. Klumperman, J., and Raposo, G. (2014). The complex ultrastructure of the endolysosomal system. Cold Spring Harb Perspect Biol 6, a016857. 10.1101/cshperspect.a016857.

23. Hoffman, D.P., Shtengel, G., Xu, C.S., Campbell, K.R., Freeman, M., Wang, L., Milkie, D.E., Pasolli, H.A., Iyer, N., Bogovic, J.A., et al. (2020). Correlative three-dimensional superresolution and block-face electron microscopy of whole vitreously frozen cells. Science 367. 10.1126/science.aaz5357.

24. Voeltz, G.K., Prinz, W.A., Shibata, Y., Rist, J.M., and Rapoport, T.A. (2006). A class of membrane proteins shaping the tubular endoplasmic reticulum. Cell 124, 573–586. 10.1016/j.cell.2005.11.047.

25. Muschalik, N., and Munro, S. (2018). Golgins. Curr Biol 28, R374–R376. 10.1016/j.cub.2018.01.006.

26. Gruenberg, J., Griffiths, G., and Howell, K.E. (1989). Characterization of the early endosome and putative endocytic carrier vesicles in vivo and with an assay of vesicle fusion in vitro. J Cell Biol 108, 1301–1316. 10.1083/jcb.108.4.1301.

27. Gopaldass, N., De Leo, M.G., Courtellemont, T., Mercier, V., Bissig, C., Roux, A., and Mayer, A. (2023). Retromer oligomerization drives SNX-BAR coat assembly and membrane constriction. EMBO J 42, e112287. 10.15252/embj.2022112287.

28. Pinot, M., Goud, B., and Manneville, J.B. (2010). Physical aspects of COPI vesicle formation. Molecular membrane biology 27, 428–442. 10.3109/09687688.2010.510485.

29. Okamoto, M., Kurokawa, K., Matsuura-Tokita, K., Saito, C., Hirata, R., and Nakano, A. (2012). High-curvature domains of the ER are important for the organization of ER exit sites in Saccharomyces cerevisiae. J Cell Sci 125, 3412–3420. 10.1242/jcs.100065.

30. Melero, A., Boulanger, J., Kukulski, W., and Miller, E.A. (2022). Ultrastructure of COPII vesicle formation in yeast characterized by correlative light and electron microscopy. Mol Biol Cell 33, ar122. 10.1091/mbc.E22-03-0103.

31. Kjeken, R., Egeberg, M., Habermann, A., Kuehnel, M., Peyron, P., Floetenmeyer, M., Walther, P., Jahraus, A., Defacque, H., Kuznetsov, S.A., and Griffiths, G. (2004). Fusion between phagosomes, early and late endosomes: a role for actin in fusion between late, but not early endocytic organelles. Mol Biol Cell 15, 345–358. 10.1091/mbc.e03-05-0334.

32. Duleh, S.N., and Welch, M.D. (2010). WASH and the Arp2/3 complex regulate endosome shape and trafficking. Cytoskeleton (Hoboken) 67, 193–206. 10.1002/cm.20437.

33. Chang, F.S., Stefan, C.J., and Blumer, K.J. (2003). A WASp homolog powers actin polymerization-dependent motility of endosomes in vivo. Curr Biol 13, 455–463. 10.1016/s0960-9822(03)00131-3.

34. Chang, F.S., Han, G.S., Carman, G.M., and Blumer, K.J. (2005). A WASp-binding type II phosphatidylinositol 4-kinase required for actin polymerization-driven endosome motility. J Cell Biol 71, 133–142. 10.1083/jcb.200501086.

35. Anitei, M., Wassmer, T., Stange, C., and Hoflack, B. (2010). Bidirectional transport between the trans-Golgi network and the endosomal system. Molecular membrane biology 27, 443–456. 10.3109/09687688.2010.522601.

36. Carreno, S., Engqvist-Goldstein, A.E., Zhang, C.X., McDonald, K.L., and Drubin, D.G. (2004). Actin dynamics coupled to clathrin-coated vesicle formation at the trans-Golgi network. J Cell Biol 165, 781–788. 10.1083/jcb.200403120.

37. Schindelin, J., Arganda-Carreras, I., Frise, E., Kaynig, V., Longair, M., Pietzsch, T., Preibisch, S., Rueden, C., Saalfeld, S., Schmid, B., et al. (2012). Fiji: an open-source platform for biological-image analysis. Nat Methods 9, 676–682. 10.1038/nmeth.2019.

38. Bolte, S., and Cordelieres, F.P. (2006). A guided tour into subcellular colocalization analysis in light microscopy. J Microsc 224, 213–232. 10.1111/j.1365-2818.2006.01706.x.

39. Ollion, J., Cochennec, J., Loll, F., Escude, C., and Boudier, T. (2013). TANGO: a generic tool for high-throughput 3D image analysis for studying nuclear organization. Bioinformatics 29, 1840–1841. 10.1093/bioinformatics/btt276.

40. Kukulski, W., Schorb, M., Welsch, S., Picco, A., Kaksonen, M., and Briggs, J.A. (2012). Precise, correlated fluorescence microscopy and electron tomography of lowicryl sections using fluorescent fiducial markers. Methods Cell Biol 111, 235–257. 10.1016/B978-0-12-416026-2.00013-3.

41. Mastronarde, D.N. (2005). Automated electron microscope tomography using robust prediction of specimen movements. J Struct Biol 152, 36–51. 10.1016/j.jsb.2005.07.007.

42. Mastronarde, D.N., and Held, S.R. (2017). Automated tilt series alignment and tomographic reconstruction in IMOD. J Struct Biol 197, 102–113. 10.1016/j.jsb.2016.07.011.

43. Kremer, J.R., Mastronarde, D.N., and McIntosh, J.R. (1996). Computer visualization of threedimensional image data using IMOD. J Struct Biol 116, 71–76. 10.1006/jsbi.1996.0013.

44. de Chaumont, F., Dallongeville, S., Chenouard, N., Herve, N., Pop, S., Provoost, T., Meas-Yedid, V., Pankajakshan, P., Lecomte, T., Le Montagner, Y., et al. (2012). Icy: an open bioimage informatics platform for extended reproducible research. Nat Methods 9, 690–696. 10.1038/nmeth.2075.

45. Paul-Gilloteaux, P., Heiligenstein, X., Belle, M., Domart, M.C., Larijani, B., Collinson, L., Raposo, G., and Salamero, J. (2017). eC-CLEM: flexible multidimensional registration software for correlative microscopies. Nat Methods 14, 102–103. 10.1038/nmeth.4170.

